# Flow cytometry-based evaluation of hepatic infection by non-fluorescent *Plasmodium* parasites

**DOI:** 10.1101/2025.07.23.666295

**Authors:** Bárbara Teixeira, Helena Nunes-Cabaço, Maria M. Mota, Diana Fontinha, Miguel Prudêncio

## Abstract

The complex life cycle of *Plasmodium* parasites, involving both liver and blood stages of infection in the mammalian host, presents significant challenges for malaria research. Although advances have been made in malaria vaccination and treatment strategies, important gaps in our understanding of the asymptomatic liver stage of *Plasmodium* infection remain. While reporter gene-expressing parasites are commonly used for drug screening and parasite biology studies during this phase of the *Plasmodium* life cycle, tools for assessing and quantifying hepatic infection in the absence of parasite-encoded reporter genes are limited. Here, we present a novel flow cytometry-based method that enables the quantitative assessment of infection of hepatic cells by non-fluorescent *Plasmodium* parasites. This method uses two parasite proteins, heat shock protein 70 (HSP70), found in the parasite cytoplasm, and upregulated in infectious sporozoites 4 (UIS4), located on the parasitophorous vacuole membrane, as markers for parasite detection and quantification. We demonstrate that the use of these markers facilitates the rapid and cost-effective quantification of hepatic infection and intracellular development of *Plasmodium* parasites devoid of fluorescent reporter genes. This method addresses critical regulatory and technical challenges to the evaluation of reporter-free whole-sporozoite vaccine candidates and could serve as a versatile tool for broader malaria research.

**Author Summary:** *Plasmodium* parasites, the causative agents of malaria, initially infect their mammalian host’s liver, where they replicate silently before entering the bloodstream and triggering disease. The hepatic stage of infection is a critical target for vaccine and drug development, but remains technically challenging to study, particularly when using parasite lines that do not express fluorescent or luminescent reporter genes. Reporter-free parasite lines are often required for regulatory reasons, particularly in the context of whole-sporozoite vaccine research. To address this limitation, we developed a flow cytometry-based method that enables the detection and quantification of hepatic infection by reporter-free *Plasmodium* parasites. The approach relies on the detection of two parasite proteins, HSP70 and UIS4, enabling the quantification of infected cells and the assessment of intracellular parasite development. This method is rapid, scalable, and cost-effective, and can be applied to *Plasmodium* lines relevant for vaccine studies. By facilitating the analysis of hepatic infection in the absence of reporter genes, our approach expands the experimental toolkit available for malaria research and supports ongoing efforts to develop interventions that target this clinically silent but biologically essential stage of the parasite’s life cycle.

## Introduction

Mammalian infection by *Plasmodium* parasites begins when an infected female *Anopheles* mosquito injects sporozoites into the host’s skin during a blood meal. These sporozoites then travel to the liver, where they invade hepatocytes and undergo asexual replication, developing into exoerythrocytic forms (EEFs) and producing thousands of merozoites (1,2). Once released into the bloodstream, merozoites infect red blood cells (RBCs) and initiate cycles of erythrocytic replication, leading to the clinical manifestations of disease and enabling transmission to the invertebrate host following ingestion of gametocytes by a mosquito during a subsequent blood meal (3,4).

Although important advances in the prevention and treatment of malaria have been made over the last few decades, this disease remains a global health challenge, with 263 million new cases and 597 000 deaths reported in 2023 (5). Moreover, while significant progress has been made in investigating the blood stage of *Plasmodium* infection, knowledge of the asymptomatic liver stage, which obligatorily precedes the onset of symptoms, remains limited (2). Nevertheless, this early stage of mammalian infection offers a critical window for intervention as it represents a bottleneck in the parasite’s life cycle and thus a key target for vaccination efforts (6).

To address this gap, a range of host cell and parasite models have been developed to study hepatic infection by *Plasmodium* and identify new points of intervention (7). Among these, rodent malaria parasites expressing different reporter genes have been successfully used to quantify infection of human hepatoma cell lines *in vitro* (8–10), both for drug screening purposes (9,11–13) and to assess the infectivity of various transgenic parasite lines (14–16). Specifically, fluorescent reporters facilitate the precise quantification of both the host cell invasion and intra-hepatic development phases of hepatic parasites by flow cytometry, making them invaluable tools in preclinical research (10,17,18).

Such models have also played a critical role in the development and evaluation of malaria vaccines targeting the liver stage. Recent advances in malaria vaccination have led to the endorsement of RTS,S/AS01 and R21/Matrix-M by the World Health Organization (WHO), two subunit vaccines that target the circumsporozoite protein (CSP), expressed on the surface of *P. falciparum* (*Pf*) sporozoites and EEFs (19,20). However, their efficacy is limited and dependent on a single antigen, which has prompted continued research into alternative approaches that could provide more robust and enduring immunity. These efforts include whole-sporozoite (WSpz) malaria vaccines, which employ live sporozoites attenuated by irradiation (radiation-attenuated sporozoites, RAS), genetic modification (genetically-attenuated parasites, GAP) or chemoprophylaxis (chemoprophylaxis and sporozoites, CPS), as the immunizing agents (reviewed in (21,22)). More recently, this list has been expanded to include PbVac, genetically engineered rodent *P. berghei* (*Pb*) parasites, used as a platform to express selected antigens from human-infective *Plasmodium* species. This strategy enables targeted antigen presentation to the human immune system within a “naturally attenuated” and cross-species immunogenic WSpz context (23,24). Besides their use as WSpz immunization agents in and of themselves, rodent *Plasmodium* parasites have played a critical role in the WSpz vaccination field. In fact, *Pb* and *P. yoelii* (*Py*) models have not only been essential for identifying target genes for deletion in *Pf*, towards the generation of *Pf*-GAP (25–28), but have also enabled the preclinical assessment and validation of *Pf*-RAS (29–31) and *Pf*-CPS (31–33) immunization strategies. However, strict regulatory requirements aimed at ensuring the safety of vaccines intended for human use demand that WSpz vaccine candidates must be devoid of selectable resistance markers or reporter genes, in order to mitigate risks associated with the introduction of foreign genetic elements (24,34). Thus, whether as vaccine candidates or as surrogates of *Pf*-based WSpz vaccines, the absence of fluorescence reporters in rodent *Plasmodium* parasites limits the use of flow cytometry as a tool for the *in vitro* assessment of their hepatic infectivity, a key aspect of their pre-clinical validation. While reverse transcription quantitative PCR (RT-qPCR) and immunofluorescence microscopy are valid alternatives to assess hepatic infection by non-fluorescent parasites, they are time-consuming, costly, and impractical for assessing multiple candidates simultaneously (17). To overcome this limitation, a novel flow cytometry-based method is presented, leveraging specific parasite proteins expressed by both sporozoites and liver stage forms to enable not only the quantification of infected cells but also the assessment of the extent of parasite development within those cells. Specifically, heat shock protein 70 (HSP70), which is primarily located in the parasite cytoplasm, where it plays a key role in cellular proteostasis (35–37), and upregulated in infectious sporozoites 4 (UIS4), a protein located on the parasitophorous vacuole membrane (PVM) that is crucial for the establishment of infection by supporting PVM growth and maintenance (38,39), are employed as markers for parasite detection and quantification. The use of HSP70 and/or UIS4 facilitates the fast and cost-effective characterization of infectivity and development of non-fluorescent *Plasmodium* parasites in hepatic cells using flow cytometry. By streamlining the evaluation of WSpz vaccine candidates, this method has the potential to support broader malaria vaccine research. Although reporter gene- expressing parasites are generally preferred for applications such as drug screening and studies of parasite biology due to their straightforward detection, this method fills a critical gap when the use of reporters is not feasible. This includes scenarios where regulatory constraints apply, such as in vaccine development, or when working with reporter-free parasite strains. Thus, the method broadens our current toolkit for research into malaria interventions and may accelerate efforts towards more effective interventions.

## Results

### HSP70- and UIS4-based assessment of hepatic Plasmodium infection by flow cytometry

In order to determine whether HSP70 and UIS4 could serve as alternatives to genetically encoded GFP for parasite detection by flow cytometry, Huh7 cells were infected with 5×10^4^ GFP-expressing *P. berghei* (*Pb*GFP) sporozoites or incubated with non-infected (NI) salivary gland material as controls. Infected and control cells were fixed and permeabilized for intracellular staining with anti-GFP, anti-HSP70, and anti- UIS4 antibodies, and then analyzed by flow cytometry at 24 and 48 h post-infection (hpi). *Pb*GFP-infected live cells were also analyzed as additional controls to exclude possible signal loss due to cell fixation and staining. Our results show that the percentage of GFP⁺ events in live cells, where GFP was detected through native fluorescence, closely matched that observed in fixed cells, in which GFP was detected via immunostaining, thereby establishing that the fixation and staining steps required for detecting HSP70 and UIS4 did not impact the accuracy of GFP detection (**Supplementary** Figure 1). A combined analysis of the GFP^+^ and either the HSP70^+^ or UIS4^+^ signals in triple-stained cells identified infected cells as double-positive events at both 24 and 48 hpi, indicating that both HSP70 and UIS4 serve as effective surrogates of genetically-encoded GFP for the analysis of hepatic cell infection (**Figure 1a**). The levels of double-positive events (GFP^+^HSP70^+^, GFP^+^UIS4^+^, and HSP70^+^UIS4^+^) closely matched those of the respective single-positive populations, indicating a strong overlap between the signals and reinforcing the specificity of these markers in identifying infected cells (**Supplementary** Figure 2). Accordingly, similar percentages of GFP^+^, HSP70^+^, and UIS4^+^ events were detected at either 24 or 48 hpi (**Figure 1b**). As expected, the overall percentage of infected cells decreased from 24 to 48 hpi, as a result of cell multiplication during this period. To further confirm that HSP70 and UIS4 can be employed to quantitatively assess the infectivity of hepatic cells by the parasite, cells were infected with increasing numbers of sporozoites and analyzed by flow cytometry 48 h later. As expected, a dose-dependent increase in the percentage of infected cells was observed, as inferred by the percentage of GFP⁺ events (**Figure 1c**, green bars). Importantly, this percentage was matched by those of the HSP70⁺ and UIS4⁺ signals (**Figure 1c**), further confirming that both markers reliably and quantitatively reflect infection levels in the absence of a fluorescent reporter. It has been shown that the geometric mean fluorescence intensity (gMFI) of GFP⁺ cells constitutes a reliable measure of the intrahepatic development of parasites expressing the GFP reporter gene (17). To assess whether HSP70 and UIS4 constitute appropriate surrogates of intracellular parasite growth, Huh7 cells were infected with 5×10^4^ *Pb*GFP sporozoites and the gMFI of the GFP⁺, HSP70⁺, and UIS4⁺ signals was determined at 24 and 48 hpi. Our results show that the gMFI of both HSP70 and UIS4 signals increases over time within infected cells, consistent with the ongoing parasite multiplication (**Figure 1d**), thus confirming their use to gauge the intrahepatic development of parasites that do not express fluorescent reporter genes. Together, these results demonstrate that both HSP70 and UIS4 constitute reliable markers for flow cytometry-based assessment of hepatic infection and multiplication of *Plasmodium* parasites.

**Figure 1.**
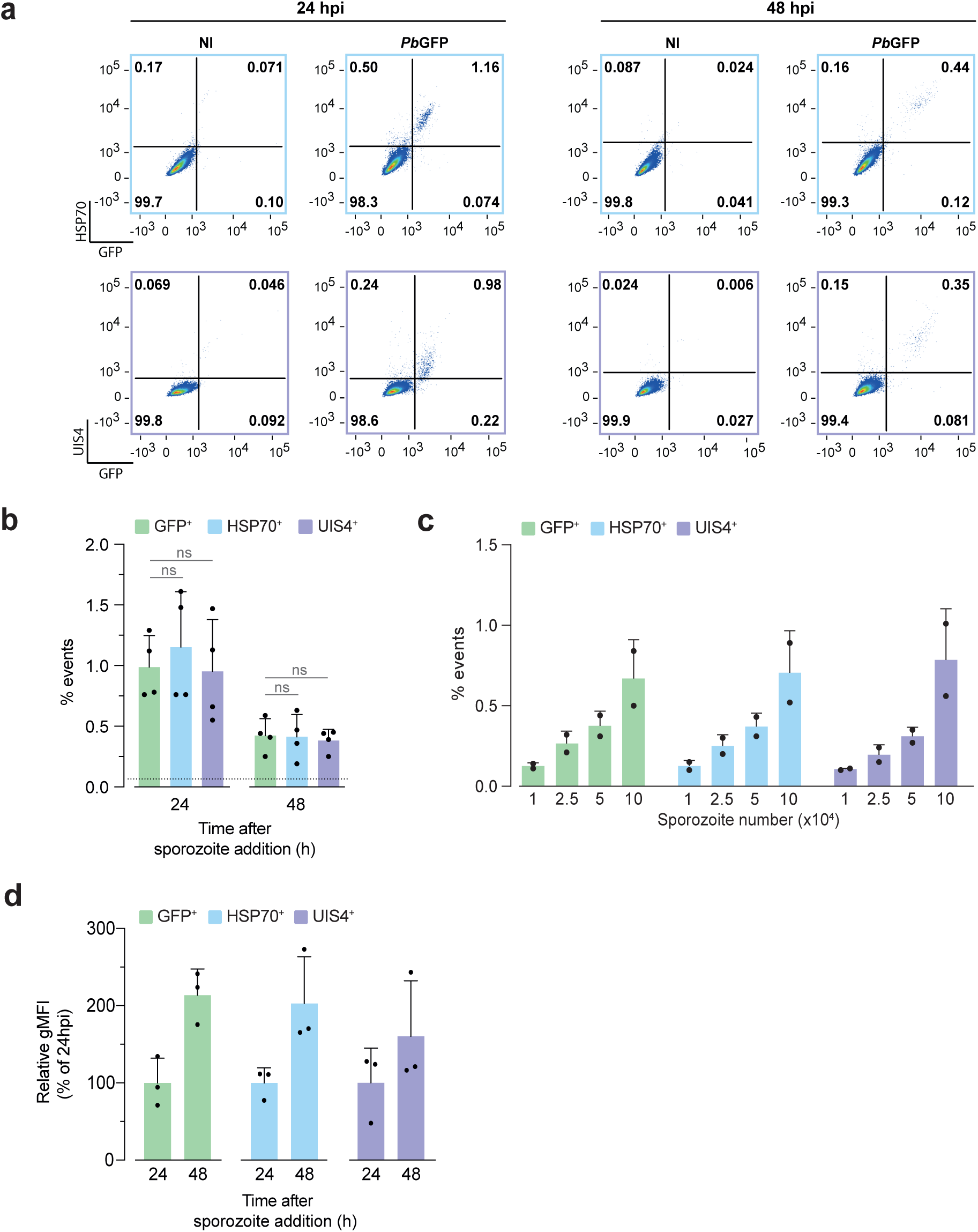
| Quantification of *Plasmodium berghei* liver stage infection and intracellular parasite development by flow cytometry. **(a)** Representative dot plots of *Pb*GFP-infected Huh7 cells at 24 and 48 hpi, stained for HSP70 (top row) and UIS4 (bottom row). The percentage of events in each quadrant is indicated. Background signal in non-infected (NI) controls was negligible. **(b)** Percentage of GFP⁺ (green), HSP70⁺ (blue), and UIS4⁺ (purple) events at 24 and 48 h after addition of 5×10^4^ *Pb*GFP sporozoites. The dashed line represents the background signal in NI controls. Results are shown as the mean ± SD of 4 independent experiments. Statistical differences relative to GFP were assessed by Kruskal-Wallis test with Dunn’s multiple comparison post-test (ns: not significant). **(c)** Percentage of GFP⁺, HSP70⁺, and UIS4⁺ events at 48 h after addition of increasing numbers of sporozoites. Results are shown as the mean ± SD of 2 independent experiments. **(d)** Relative geometric mean fluorescence intensity (gMFI) of GFP, HSP70, and UIS4 signals at 24 and 48 hpi. gMFI values correspond to the geometric mean fluorescence intensity for each marker, normalized to the respective values at 24 hpi. Results are shown as the mean ± SD of 3 independent experiments; dots represent individual replicates.

### Flow cytometry-based assessment of hepatic infection by reporter-free whole-sporozoite vaccine candidates

To validate the use of flow cytometry for quantification of infection by parasite lines that do not carry reporter genes, particularly those employed in WSpz vaccine research, Huh7 cells were infected with 5×10^4^ non-attenuated (*Pb*WT) or radiation-attenuated (*Pb*RAS) wild-type sporozoites, and early (*Pb*EA-GAP) and late (*Pb*LA- GAP) arresting GAP sporozoites. *Pb*RAS and *Pb*EA-GAP parasites arrest early after hepatocyte invasion, typically within the first 12-24 h (25,40), while *Pb*LA-GAP parasites arrest at later stages, often after several rounds of replication but before completion of liver stage development (41). Cells were analyzed by flow cytometry 48 h later. Our results show a clear correlation between the intensity of the gMFI of both the HSP70 (**Figure 2a**) and the UIS4 (**Figure 2b**) signals, and the expected extent of intrahepatic parasite development, indicating that both markers constitute appropriate surrogates of parasite development within Huh7 cells. Notably, similar results were obtained when employing HepG2 cells, underscoring the robustness of the method across different host cell lines (**Supplementary** Figure 3). To further confirm that HSP70 and UIS4 signal intensities constitute appropriate surrogates of intrahepatic parasite growth *in vitro*, we assessed both the total parasite load, as measured by RT-qPCR (**Figures 2c,d**), and parasite size, determined by immunofluorescence microscopy (**Figures 2e,f**) in samples obtained in parallel with those employed for flow cytometry analysis. Our results showed a clear correlation between the gMFIs of both the HSP70 and the UIS4 signals and either the hepatic parasite load (**Figure 2c,d**, **Supplementary** Figure 4a) or the EEF size (**Figure 2e,f, Supplementary** Figure 4b) in those samples. Collectively, these results affirm the suitability of HSP70 and UIS4 to assess the development of fluorescent reporter-free parasites by flow cytometry.

**Figure 2.**
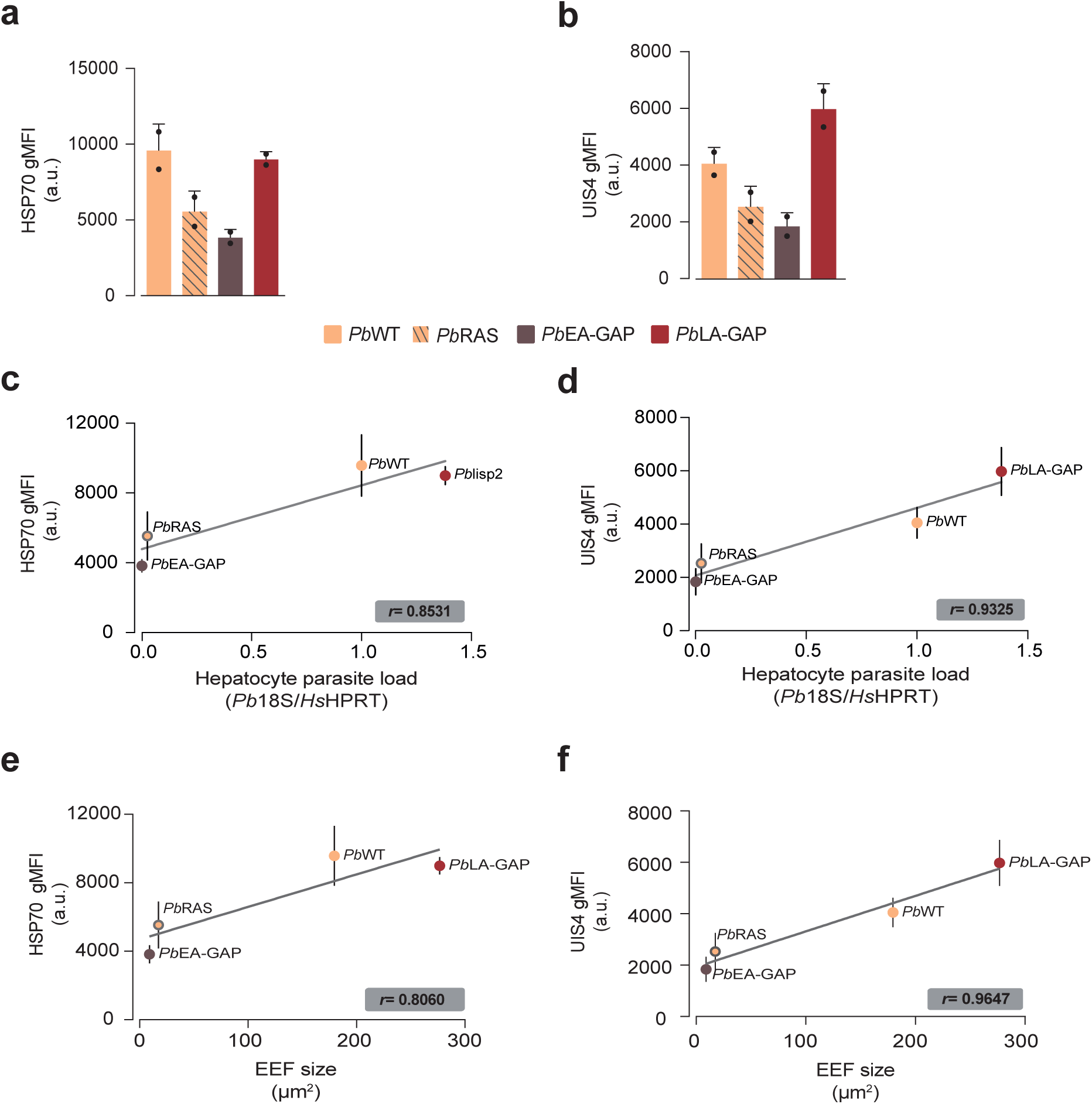
| Flow cytometry enables detection and developmental assessment of non-fluorescent *Plasmodium* liver stage parasites using HSP70 and UIS4 Geometric mean fluorescence intensity (gMFI) in arbitrary units (a.u.) of **(a)** HSP70 and **(b)** UIS4 in Huh7 cells infected with 5×10^4^ *Pb*WT, *Pb*RAS, *Pb*EA-GAP, or *Pb*LA- GAP sporozoites, at 48 h post-infection (hpi). gMFI values correspond to the geometric mean fluorescence intensity for each marker. Results are shown as the mean ± SD from 2 independent experiments. Correlation between **(c)** HSP70 and **(d)** UIS4 gMFI values measured by flow cytometry and parasite load determined by RT-qPCR targeting *Pb18s* rRNA. RT-qPCR data are represented as fold change relative to cells infected with *Pb*WT parasites. Correlation between **(e)** HSP70 and **(f)** UIS4 gMFI values and exoerythrocytic form (EEF) size determined by microscopy using the UIS4 signal (in µm²). Linear regression was used for visualization, and *r* values represent Pearson correlation coefficients.

## Discussion

The liver stage of *Plasmodium* infection remains a critical yet under-addressed window in malaria research, particularly in settings where reporter-free parasites are required. Here, we provide a practical and scalable solution to this longstanding technical bottleneck. To that end, we developed and validated a flow cytometry-based method to quantify *Plasmodium* liver stage infection using HSP70 and UIS4 as intracellular markers. Both proteins reliably identified infected cells, with signal intensity correlating with parasite development. Building on this marker combination, the method enabled quantification of infection across a range of input sporozoite doses, was applicable to non-reporter parasite lines, and showed strong agreement with established infection metrics such as RT-qPCR and immunofluorescence microscopy. While RT-qPCR and immunofluorescence microscopy have long served as the gold standards for assessing liver stage infection, their low throughput and technical demands significantly limit scalability. Our method circumvents these limitations by enabling parasite developmental staging of individual infected cells, as well as high-throughput compatibility in a single assay, all without the need for reporter gene. This is particularly relevant for WSpz vaccine development, where the use of genetically modified reporter parasites is often prohibited (24,34). The ability to monitor infection and development in these contexts with such precision and reproducibility could streamline preclinical assessment pipelines and facilitate regulatory compliance.

In line with this, we observed that the gMFI values reflect the extent of parasite development, as shown by comparable HSP70 gMFI between *Pb*LA-GAP- and *Pb*WT- infected cells at 48 hpi. This is consistent with previous reports showing that *Pb*LA- GAP parasites arrest around 60 hpi *in vivo*, and possibly slightly later *in vitro* (41), supporting the utility of gMFI as a proxy for intrahepatic parasite growth. Interestingly, the UIS4 gMFI was higher in cells infected with *Pb*LA-GAP than with *Pb*WT, consistent with observations that this parasite line can form larger, albeit non-productive, liver stage forms *in vitro* (41,42), potentially leading to increased accumulation of UIS4 protein at the PVM, where it localizes (38,39).

Given these findings at later stages of hepatic infection, we next asked whether the same markers could be used to assess early stages of infection. We tested whether the intracellular markers HSP70 and UIS4 could be used to assess hepatic cell invasion by flow cytometry at 2 hpi, as previously demonstrated for GFP-expressing parasites (17). However, we were unable to detect reliable fluorescence signal with either marker at that early time point, suggesting that these proteins are not yet expressed or accumulated to measurable levels. While surface markers, such as CSP, are commonly used to study parasite entry into hepatocytes, they do not provide information on the parasite’s intracellular development (43). As such, we focused our analyses on later time points (24 and 48 hpi), when HSP70 (42,44) and UIS4 (38) expression is more robust and better reflects parasite maturation within host cells.

These observations highlight the need for tools that can quantify both infection and developmental progression at the liver stage, especially in experimental contexts where reporter genes are absent due to technical or regulatory constraints. Luminescence-based assays are commonly used in liver stage *Plasmodium* studies and provide a sensitive, global measure of parasite burden (7,9). However, they do not offer single-cell resolution and cannot distinguish whether changes in signal result from differences in invasion efficiency or subsequent parasite development, which may complicate data interpretation (7). To gain spatial resolution and characterize developmental arrest, immunofluorescence microscopy has been widely employed to characterize GAPs developed as WSpz vaccine candidates. Parasites lacking *uis3* (38) and *slarp* (45) were shown to arrest early in liver stage development based on microscopy analysis. Similarly, *sap1* (46) deletion mutants and *b9/slarp* double knockout (25) in *P. berghei* were characterized primarily by microscopy to confirm developmental arrest and infectivity defects. While microscopy provides a detailed visualization of parasite infection and development, it can be labour-intensive and is low throughput (7). To overcome these limitations and enable higher-throughput analysis, flow cytometry has been successfully used to quantify liver stage infection and development using *Pb*GFP-expressing parasites, providing a robust and reproducible alternative for fluorescent reporter lines (17,47,48). Building on this, our flow cytometry-based approach expands the applicability of this technique by enabling the assessment of liver stage infection and progression using intracellular parasite markers. This allows for quantitative, rapid, and accessible evaluation of parasites lacking reporter genes. Such advancement is particularly valuable in the context of WSpz vaccine research, where objective, scalable, and standardized tools are urgently needed. Our flow cytometry-based method allows independent assessment of infection rates (based on the quantification of HSP70- and/or UIS4-positive events) and developmental progression (based on the gMFI values of these markers), enabling a detailed characterization of parasite dynamics post-infection. An additional advantage is the accessibility of the UIS4 antibody, which is commercially available and has been widely employed in multiple studies (49,50), making the method easier to adopt.

A limitation of our approach is its current validation in hepatoma cell lines, which may not fully recapitulate the microenvironment of primary hepatocytes or *in vivo* liver tissue. Nonetheless, the method’s adaptability and compatibility with primary cells, as previously used in PbVac studies (23,24), support its future deployment in more physiologically relevant systems Flow cytometry further allows for the inclusion of additional markers in the antibody panel, such as other parasite proteins and host cell surface or intracellular proteins whose expression might be altered by infection, enabling multiparametric analysis of host-parasite interactions. This represents a key advantage over microscopy, which is limited by the number of fluorophores that can be simultaneously visualized. Several transcriptomic studies have identified host proteins whose expression is modulated during *Plasmodium* liver stage infection, including components of stress response and metabolic pathways (51–54). In parallel, parasite proteins expressed at distinct stages of liver stage development represent additional targets that could be incorporated into future flow cytometry analyses (55–57). Expanding this method to liver stage models of human-infecting species, such as *P. falciparum*, could also significantly broaden its translational relevance. Given the pivotal role of the liver stage of infection as both the first step in disease progression and a potential bottleneck for transmission, accessible and quantitative methods for its evaluation remain a priority (7). Our method directly addresses this need by offering a flexible and informative platform for investigating hepatic infection. As such, it holds promise for diverse applications, from vaccine candidate evaluation to fundamental studies of host-parasite interactions. By enabling quantitative, scalable, and reporter-independent analysis of hepatic infection, this method not only expands the experimental toolbox but also lays the groundwork for accelerating discovery across both fundamental research and translational applications in malaria.

## Materials and Methods

### Cells, parasites, and infection

Huh7 cells, a human hepatoma cell line, were cultured in Roswell Park Memorial Institute 1640 (RPMI) medium supplemented with 1% (v/v) penicillin/streptomycin, 10% (v/v) fetal bovine serum (FBS), 1% (v/v) glutamine, 1% (v/v) non-essential amino acids, and 10 mM 4-(2-hydroxyethyl)-1-piperazineethanesulfonic acid (HEPES), pH 7.0 (all from Gibco/Invitrogen)- cRPMI. HepG2 cells, another human liver-derived cell line, were cultured in Dulbecco’s Modified Eagle Medium (DMEM), supplemented with 1% (v/v) penicillin/streptomycin, 10% (v/v) FBS, and 1% (v/v) glutamine - cDMEM. Both cell lines were maintained at 37 °C in a humidified atmosphere with 5% CO_2_. Female *Anopheles stephensi* mosquitoes, reared at the Gulbenkian Institute for Molecular Medicine (Lisbon, Portugal), were infected with the following *Plasmodium berghei* parasite lines: (i) GFP-expressing *Pb*ANKA (GFP-CON; 259 cl2, mutant RMgm-5; www.pberghei.eu), referred as *Pb*GFP (10) (ii) *Pb*ANKA (cl15cy1), referred as *Pb*WT (58) (iii) *PbΔb9Δslarp* parasites (1844 cl1, mutant RMgm-1141; www.pberghei.eu), which are genetically attenuated by the deletion of the *b9* and *slarp* genes (25), referred as *Pb*EA-GAP and (iv) *PbΔmei2Δlisp2* parasites (2900 cl3, mutant RMgm-4941; www.pberghei.eu), in which the expression of MEI2 and LISP2 proteins has been abrogated (59), referred as *Pb*LA-GAP. Non-infected salivary gland material was obtained by dissection of the salivary glands of non-infected *A. stephensi* mosquitoes. Cells were plated (5 × 10^4^ per well) on 24-well plates the day before infection. Cells were infected with freshly dissected *P. berghei* sporozoites, followed by centrifugation at 600 xg for 5 min at room temperature (RT). Cells were incubated at 37 °C in a humidified atmosphere with 5% CO_2_ for 2 h, following which the medium was replaced by fresh supplemented culture medium containing 50 µg/mL Gentamycin and 0.85 µg/mL Amphotericin B (both from Gibco). Cells were incubated at 37 °C in a humidified atmosphere with 5% CO_2_ and maintained under these conditions until the selected time point post-sporozoite addition. Radiation-attenuated sporozoites were obtained by exposure of sporozoites to 16,000 rad of γ-radiation on a Gammacell® 3000 Elan irradiator (Best Theratronics).

### Flow cytometry

Cell samples were collected for flow cytometry analysis at the selected time points post-sporozoite addition. Cells were washed once with 500 µL of phosphate-buffered saline (PBS) and incubated with 150 µL of trypsin for 5 min at 37 °C. After trypsinization, 200 µL of fresh supplemented culture medium was added to neutralize the trypsin, and the cell suspension was collected to 1.5 mL microtubes. Wells were washed with an additional 100 µL of medium to ensure complete collection of cells. The cell suspension was centrifuged at 600 xg for 5 min, and the pellet was re- suspended in 100 µL of PBS before transferring the cells to a U-bottom 96-well plate. Cells were then fixed by adding 100 µL of IC fixation buffer (eBioscience, 00-8222-49) and incubating for 20 min at 4 °C. After fixation, the plate was centrifuged at 600 xg for 5 min, the supernatant was discarded by flicking the plate, and the cells were washed with 150 µL of FACS buffer (2% FBS and 2 mM EDTA in PBS). Cells were immediately stained or stored at 4°C overnight. For cell permeabilization, 100 µL of Perm buffer (1:10 dilution in distilled water of 10x Permeabilization Buffer, eBioscience, 00-8333-56) was added. The plate was then centrifuged at 600 xg for 5 min, and the buffer was removed. To block Fc receptors and prevent non-specific binding of antibodies, Human TruStain FcX Fc Receptor blocking solution (1:50 dilution in FACS buffer, Biolegend, 422301) was added for 30 min at 4 °C. Cells were then washed with 100 µL of FACS buffer. For intracellular staining, cells were incubated with 20 µL of the primary antibody, diluted in Perm buffer, for 30 min at 4 °C. After incubation, cells were washed with 100 µL of FACS buffer, centrifuged, and the supernatant was flicked out. The cells were then incubated with 20 µL of secondary antibody, diluted in Perm buffer, for 30 min at 4 °C. Following the final antibody incubation, cells were washed with 100 µL of FACS buffer. Finally, cells were re- suspended in 250 µL of FACS buffer and transferred to cytometry tubes. Cells were analyzed on a LSRFortessa^TM^ Cell Analyzer (BD Biosciences) immediately after preparation or stored at 4°C in the dark until analysis, within 12 h. All events were acquired for each sample and gated using forward scatter (FSC) versus side scatter (SSC) plots. Cells treated with non-infected salivary gland material at the same time as the addition of the sporozoites were used as controls to ensure that the gates did not contain false-positive events. Data analysis was carried out using FlowJo v10.7.1 software (Tree Star Inc.). The primary antibodies used for intracellular staining included anti-*P. berghei* HSP70 (1:50, mouse monoclonal, 2E6, produced in-house (44)), anti-*P. berghei* UIS4 (1:50, goat polyclonal, SicGen, AB0042-200), and anti-GFP polyclonal antibody Alexa Fluor 488-conjugated (1:300, Thermo Fisher Scientific, A21311). Donkey anti-mouse Alexa Fluor Plus 647 (1:500, Thermo Fisher Scientific, A32787) and donkey anti-goat R-PE-conjugated (1:200, Proteintech, SA00008-3) were employed as secondary antibodies.

### Immunofluorescence microscopy

Cells plated on glass coverslips were fixed in 4% paraformaldehyde (ChemCruz) for 10 min at RT and permeabilized/blocked with a solution of 1% BSA - 0.2% Triton X- 100 in PBS (perm/block) for 1 h at RT. For immunostaining, samples were incubated with primary antibodies diluted in perm/block solution for 1.5 h at RT, washed with PBS, incubated for 1 h at RT with the Alexa Fluor-conjugated secondary antibodies and Hoechst 33342 (1:1000, Invitrogen, H1399) diluted in perm/block and washed again with PBS. The coverslips were then mounted on microscope slides with Fluoromount G (Invitrogen). Primary antibodies used for fluorescence microscopy included anti-*P. berghei* HSP70 (1:50, mouse monoclonal, 2E6, produced in-house) and anti-*P. berghei* UIS4 (1:300, goat polyclonal, SicGen, AB0042-200). The secondary antibodies used were donkey anti-goat Alexa Fluor 568 (1:1000, Thermo Fisher Scientific, A11057) and donkey anti-mouse Alexa Fluor 647 (1:1000, Thermo Fisher Scientific, A31571). All images were acquired on Zeiss Celldiscoverer 7 with LSM 900 microscope (Zeiss) and processed using Fiji (version 1.54n) (60), ImageJ (version 1.53t) (61), and ZEN Blue Edition software (version 3.4.91.0000).

### cDNA synthesis and quantitative reverse-transcriptase PCR

Cells were washed with 500 µL of PBS and collected at the selected time points in 1 mL of TRizol reagent (Invitrogen) and immediately stored at -80 °C until further processing. Total RNA extraction from cells was performed using NZY Total RNA Isolation Kit (NZYTech), following manufacturer’s instructions, and quantified in a Nanodrop 2000 (Thermo Fischer Scientific). cDNA was synthesized from 1 µg of RNA using a mix of random hexomers (2.5 ng/µL), dNTPs (0.5 mM), ribonuclease inhibitor (1 U/µL) and reverse transcriptase (10 U/µL) in a reaction buffer for reverse transcriptase 1x (all reagents from NZYTech). The reverse transcriptase reaction was performed in a T100^TM^ Thermal Cycler (Bio-Rad) with the following conditions: 25 °C, 10 min; 55 °C, 30 min, and 85 °C, 5 min. cDNA was stored at -20 °C until further use.

RT-qPCR was performed to detect *P. berghei 18s* (*Pb18s*) RNA and *Homo sapiens hypoxanthine-guanine phosphoribosyltransferase* (*Hshprt*) RNA in the iTaq Universal SYBR Green Supermix (Bio-Rad) using RT-PCR 7500 Fast cycler (Applied Biosystems) with the following conditions: 50 °C, 2 min; 95 °C, 10 min; 40 cycles at 95 °C, 5 s and 60 °C, 1 min, 95 °C, 15 s, and 60 °C, 1 min. Gene expression values were calculated based on the ΔΔC_T_ method, using *Hsprt* as the reference gene, and the mean of the control group as the calibrator to which all the other samples were compared. The following primers were used: *Pb18s* – forward: AGGGAGCCTGAGAAATAG and reverse: GTCACTACCTCTCTTATTTAGAA; and *Hshprt* – forward: TTTGCTGACCTGCTGGATTAC and reverse: CAAGACATTCTTTCCAGTTAAAGTTG.

### Statistical analysis

Results are presented as mean ± SD. Statistical analyses were performed using GraphPad Prism 10 (GraphPad Software, Inc). All the data were compared using the Kruskal-Wallis with the Dunn’s multiple comparison post-test. Significance of the differences observed is indicated in each figure, ns: not significant.

## Acknowledgments

We thank Ana Parreira and Carla Oliveira for mosquito rearing and *Plasmodium* parasite infections, and Diana Moita and Eyob A. Workneh for support with sporozoite irradiation procedures and microscopy image acquisition. We are also grateful to the Flow Cytometry Facility at the Gulbenkian Institute for Molecular Medicine for technical assistance. BT was supported by a Fundação para a Ciência e a Tecnologia (FCT) fellowship (UI/BD/151170/2021) and GIMM/BI/16-2025. The study was supported by FCT Grant 2022.02624.PTDC to DF, by “la Caixa” Foundation Grant HR21-848 to MP, and by European Research Council (ERC) Grant ERC-2022-AdG-101097801- PASSAGE to MMM.

---

**Supplementary Figure 1.**
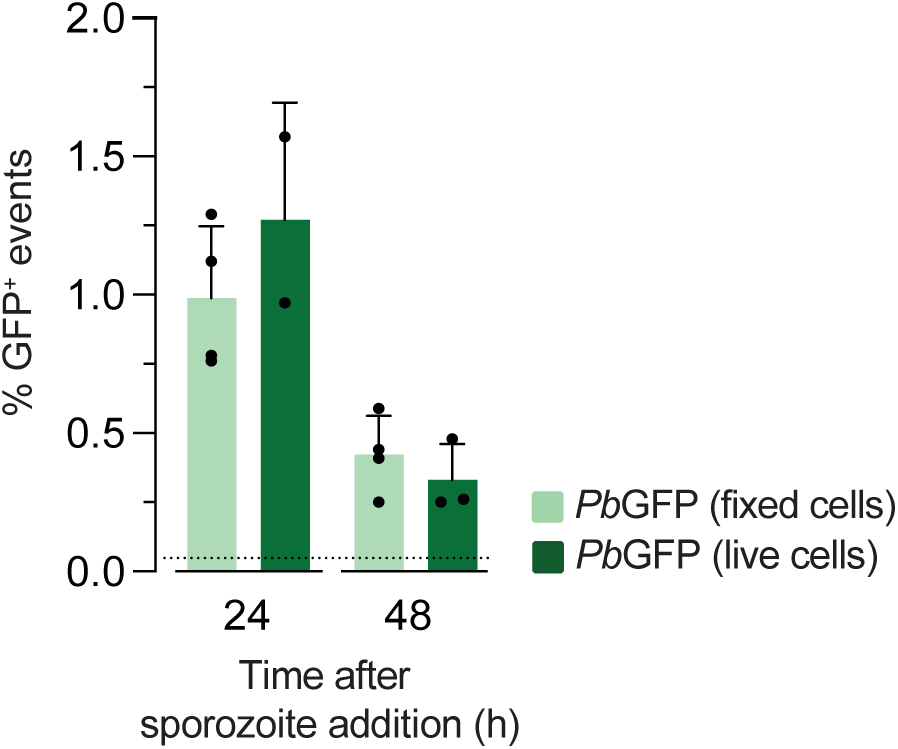
| Validation of GFP detection in fixed and stained cells. To confirm that fixation and staining did not compromise GFP detection, the percentage of GFP⁺ events was compared between fixed and stained cells and live, unfixed cells. Huh7 cells were infected with *Pb*GFP sporozoites and analyzed at 24 and 48 h post-infection (hpi). For fixed samples, GFP was detected by immunostaining, while for live samples, GFP was detected via its native fluorescence. The dashed line represents the background signal in NI controls. Results are shown as mean ± SD of 3 independent experiments.

**Supplementary Figure 2.**
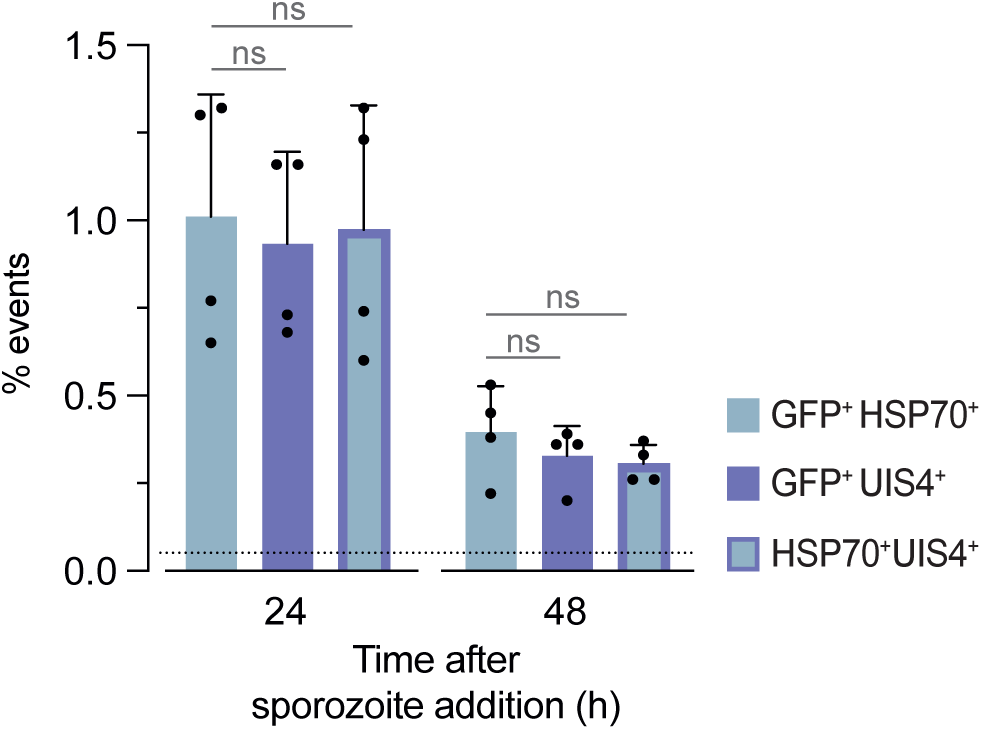
| Quantification of double-positive events supports the concordance between GFP, HSP70, and UIS4 signals. Huh7 cells were infected with 5×10^4^ GFP-expressing *P. berghei (Pb*GFP) sporozoites and stained for the parasite markers HSP70 and UIS4 at 24 and 48 h post-infection (hpi). Bar plots represent the proportion of GFP⁺HSP70⁺, GFP⁺UIS4⁺, and HSP70⁺UIS4⁺ events 24 and 48 h later. Statistically significant differences relative to GFP were assessed through the Kruskal-Wallis test with Dunn’s multiple comparison post-test (ns: not significant). Results are shown as mean ± SD of 4 independent experiments.

**Supplementary Figure 3.**
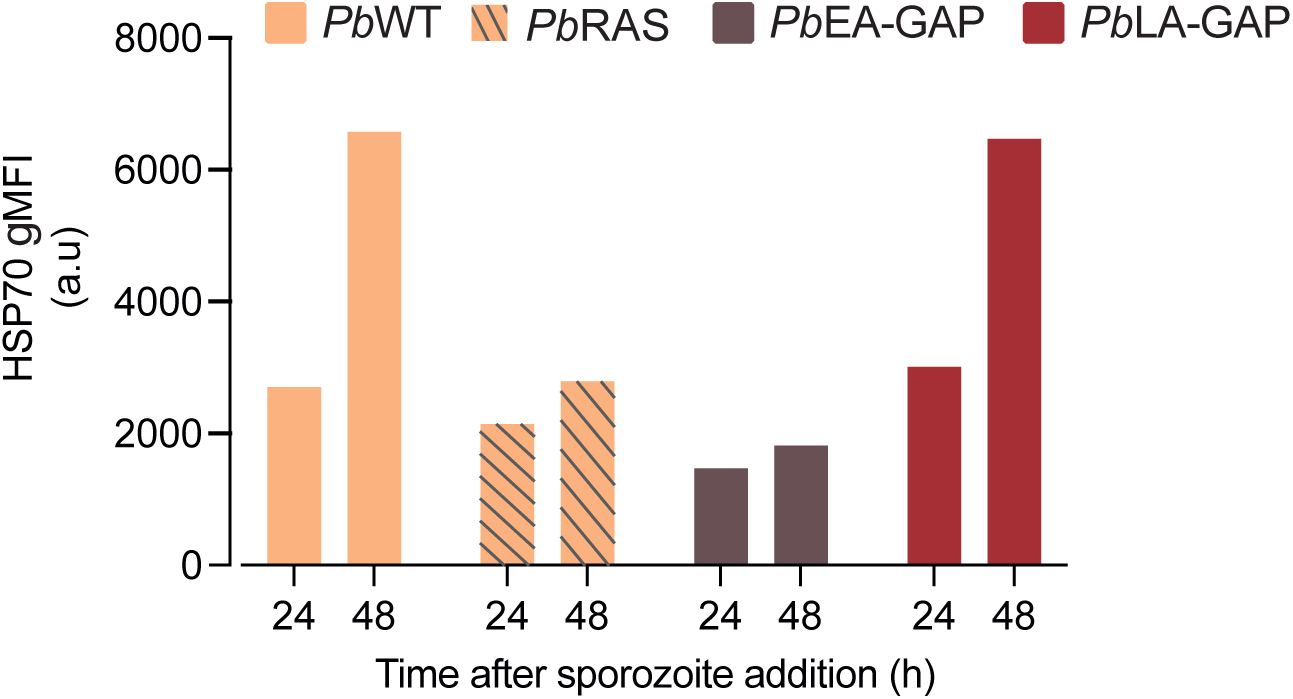
| Application of the method across hepatocyte cell lines. Geometric mean fluorescence intensity (gMFI) in arbitrary units (a.u.) of HSP70 shown for HepG2 cells infected with *P. berghei* lines that do not express fluorescent proteins (*Pb*WT, *Pb*RAS, *Pb*EA-GAP, or *Pb*LA-GAP) at 24 and 48 h post-infection (hpi). gMFI values correspond to the geometric mean fluorescence intensity for each marker. These results correspond to a single experiment.

**Supplementary Figure 4.**
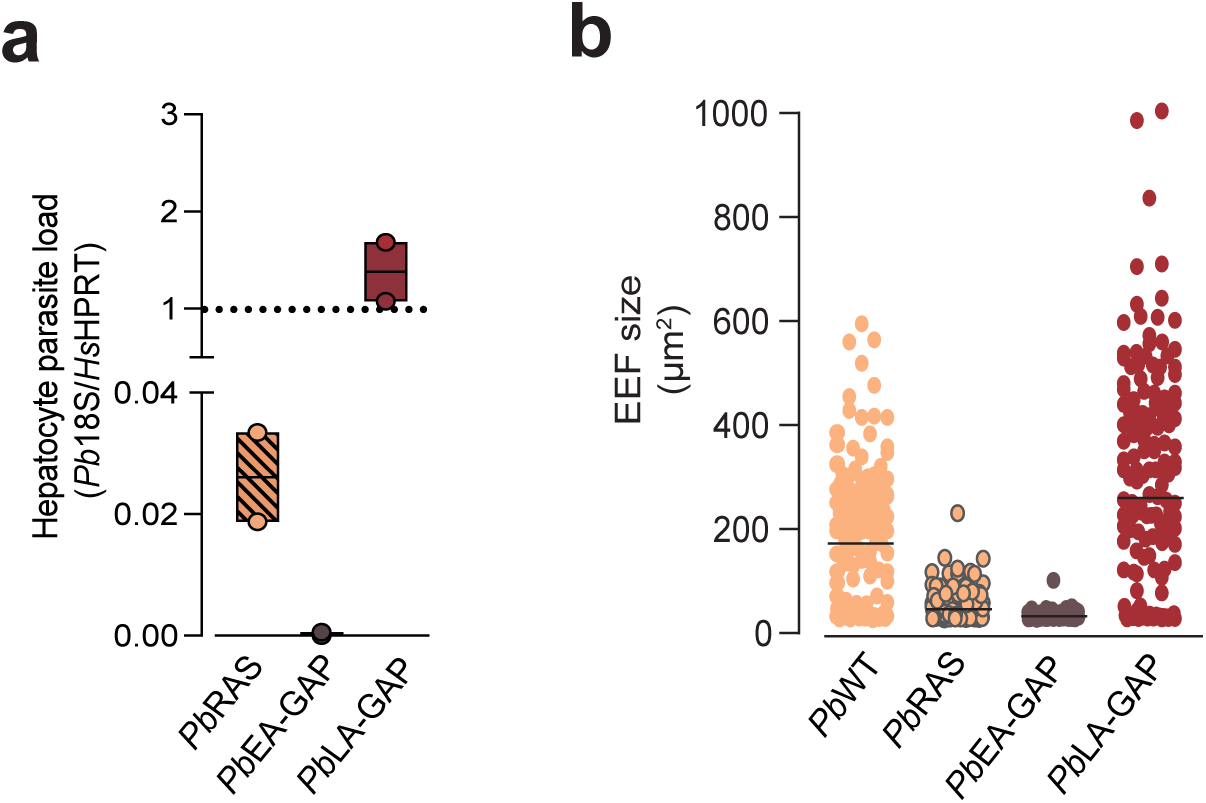
| Additional validation of the flow cytometry-based method. **(a)** RT-qPCR data used for the correlation analysis presented in Figure 2, represented as fold change relative to cells infected with *Pb*WT parasites (dashed line). Results are presented as boxplots (min-to-max) with a line indicating the mean, based on 2 independent experiments. **(b)** Microscopy data used to compute the correlation analysis. The size of exoerythrocytic forms (EEF) is shown in µm². Results are shown as dot plots with a line representing the mean, based on 2 independent experiments.

